# Influenza Virus Infection Model With Density Dependence Supports Biphasic Viral Decay

**DOI:** 10.1101/253401

**Authors:** Amanda P. Smith, David J. Moquin, Veronika Bernhauerova, Amber M. Smith

## Abstract

Mathematical models that describe infection kinetics help elucidate the time scales, effectiveness, and mechanisms underlying viral growth and infection resolution. For influenza A virus (IAV) infections, the standard viral kinetic model has been used to investigate the effect of different IAV proteins, immune mechanisms, antiviral actions, and bacterial coinfection, among others. We sought to further define the kinetics of IAV infections by infecting mice with influenza A/PR8 and measuring viral loads with high frequency and precision over the course of infection. The data highlighted dynamics that were not previously noted, including viral titers that remain elevated for several days during mid-infection and a sharp 4-5 log_10_ decline in virus within one day as the infection resolves. The standard viral kinetic model, which has been widely used within the field, could not capture these dynamics. Thus, we developed a new model that could simultaneously quantify the different phases of viral growth and decay with high accuracy. The model suggests that the slow and fast phases of virus decay are due to the infected cell clearance rate changing as the density of infected cells changes. To characterize this model, we fit the model to the viral load data, examined the parameter behavior, and connected the results and parameters to linear regression estimates. The resulting parameters and model dynamics revealed that the rate of viral clearance during resolution occurs 25 times faster than the clearance during mid-infection and that small decreases to this rate can significantly prolong the infection. This likely reflects the high efficiency of the adaptive immune response. The new model provides a well-characterized representation of IAV infection dynamics, is useful for analyzing and interpreting viral load dynamics in the absence of immunological data, and gives further insight into the regulation of viral control.

## Introduction

Influenza A virus (IAV) is a leading cause of lower respiratory tract infections and causes a significant amount of morbidity and mortality [28, 39, 49], with over 15 million individuals infected and more than 200,000 hospitalizations each year in the U.S. [50]. Vaccination against influenza viruses remains the most effective measure to prevent infection, but the large number of antigenically distinct strains, the emergence of new strains, and the low efficacy of antivirals make combatting the disease challenging. New therapeutic strategies are thus necessary and may require modulation of different viral control mechanisms, which are not entirely understood for IAV infection. Thus, it is critical to gain a deeper understanding of the infection kinetics, including determining the time scales, magnitudes, contribution, and interrelatedness of different control processes throughout IAV infection.

Kinetic modeling of *in vivo* infection processes provides important insight into viral growth and decay, host immune responses, antiviral actions, and multi-pathogen interactions. Remarkably, as few as 3-4 equations for target cells, infected cells, and virus can accurately describe viral load dynamics for a variety of virus infections (e.g., IAV, HIV, HCV, Zika virus, and West Nile Virus [1, 2, 5, 31, 34]). For IAV infections, numerous studies have used these simple models with great success to elucidate mechanisms during IAV infection and during IAV coinfection with bacterial pathogens (reviewed in [3, 7, 41–43]). However, investigating mechanisms of immune control is often inhibited by insufficient data, which limits effective model calibration and selection. Further, it can be difficult to distinguish between mechanisms because a viral kinetic model that excludes equations and terms for specific immune responses can fit viral load dynamics with ease.

To aid interpretation of model results and gain insight into the mechanisms of infection, previous studies have used linear regression and approximate solutions to the viral kinetic model (derived by [45]) to identify how different processes (e.g., virus infection, production, and clearance) contribute to viral load dynamics throughout the course of infection [5, 16, 18, 19, 22, 26, 29, 32, 36, 40, 45, 46]. In the initial hours of infection, virus quickly infects cells or is cleared. Following an eclipse phase, virus production begins and virus increases exponentially for ~2 d. This initial growth can be approximated by a linear function of the log_10_ viral titers or by *V*(*t*) = *e*^λ*t*^, where λ is a combination of all infection processes and is equivalent to the log-linear slope [45, 46]. After this growth phase, virus peaks and begins to decline until the infection is resolved. Virus decay is typically exponential in nature and can be approximated in a similar fashion as the growth phase. That is, *V*(*t*) = *e*^−δ*t*^, where δ is the infected cell death rate and the sole process dictating the viral decay dynamics. Here, the log-linear slope is an estimate of the infected cell death rate [45, 46].

Although these dynamics and approximations have improved our knowledge of viral kinetics, some dynamical features, such as the plateauing of virus following the peak (reviewed in [42]) cannot be explained by current kinetic models that exclude equations for immune factors. One model could reproduce the plateauing of virus through modeling interferon and an interferon-induced adaptive immune response [33]. The study concluded that specific equations for the innate and adaptive responses were necessary. However, quantitative immunological data was not used to support model selection, parameterization, and conclusions. This type of data is scarce and has been a limiting factor of modeling studies. With viral loads as the most prevalent type of data, models that limit the number of parameters and equations remain desirable. However, even most viral load data is insufficiently quantitative to confidently detect features like a mid-infection plateau and build appropriate mathematical models.

Here, we first sought to increase the quality and quantity of viral load data in order to improve predictive power of mathematical models and gain a deeper insight into the kinetics of viral resolution. To do this, we measured viral loads daily from groups of BALB/cJ mice infected with influenza A/Puerto Rico/8/34 (H1N1) (PR8). In addition, we tightly controlled the experimental conditions and repeated the experiment numerous times to ensure reproducibility and identify data with meaningful biological heterogeneity (i.e., due to an underlying mechanism) versus data with experimental heterogeneity (i.e., due to poor technique). The high resolution of these data defined important dynamical features, including a long plateau phase followed by a rapid decay phase. Because current viral kinetic models cannot reproduce these data, we developed a new model that incorporated a density-dependent decay of infected cells and could accurately describe the observed viral load dynamics. We used a rigorous fitting scheme to estimate the model parameters and infer important dynamics. Subsequent linear regression analysis and sensitivity analysis aided effective interpretation of the model results and direct comparison with published results. The data, model, and analyses provide a robust quantification of IAV infection kinetics and indicate that the rate of virus clearance changes with respect to the density of infected cells.

## Materials and Methods

### Use of Experimental Animals

All experimental procedures were approved by the Animal Care and Use Committee at SJCRH under relevant institutional and American Veterinary Medical Association guidelines and were performed in a Biosafety level 2 facility that is accredited by AALAAS.

### Mice

Adult (6 week old) female BALB/cJ mice were obtained from Jackson Laboratories (Bar Harbor, ME). Mice were housed in groups of 5 mice in high-temperature 31.2cm × 23.5cm × 15.2cm polycarbonate cages with isolator lids. Rooms used for housing mice were maintained on a 12:1 2-hour light:dark cycle at 22 ± 2°C with 50% humidity in the biosafety level 2 facility at St. Jude Children’s Research Hospital (Memphis, TN). Prior to inclusion in the experiments, mice were allowed at least 7 days to acclimate to the animal facility such that they were 7 weeks old at the time of infection. Laboratory Autoclavable Rodent Diet (PMI Nutrition International, St. Louis, MO) and autoclaved water were available ad libitum. All experiments were performed under an approved protocol and in accordance with the guidelines set forth by the Animal Care and Use Committee at St. Jude Children’s Research Hospital.

### Infectious Agents

All experiments were done using the mouse adapted influenza A/Puerto Rico/8/34 (H1N1) (PR8).

### Infection Experiments

The viral infectious dose (TCID_50_) was determined by interpolation using the method of Reed and Muench [37] using serial dilutions of virus on Madin-Darby canine kidney (MDCK) cells. Mice were intranasally inoculated with 75 TCID_50_ PR8 in 100 μl Mice were weighed at the onset of infection and each subsequent day for illness and mortality. Mice were euthanized if they became moribund or lost 30% of their starting body weight. Each experiment was repeated numerous times to ensure reproducibility. Two complete experiments (10 animals per time point) were used for these studies. The raw data is shown in Figure 1A and is available upon request.

**Figure 1.**
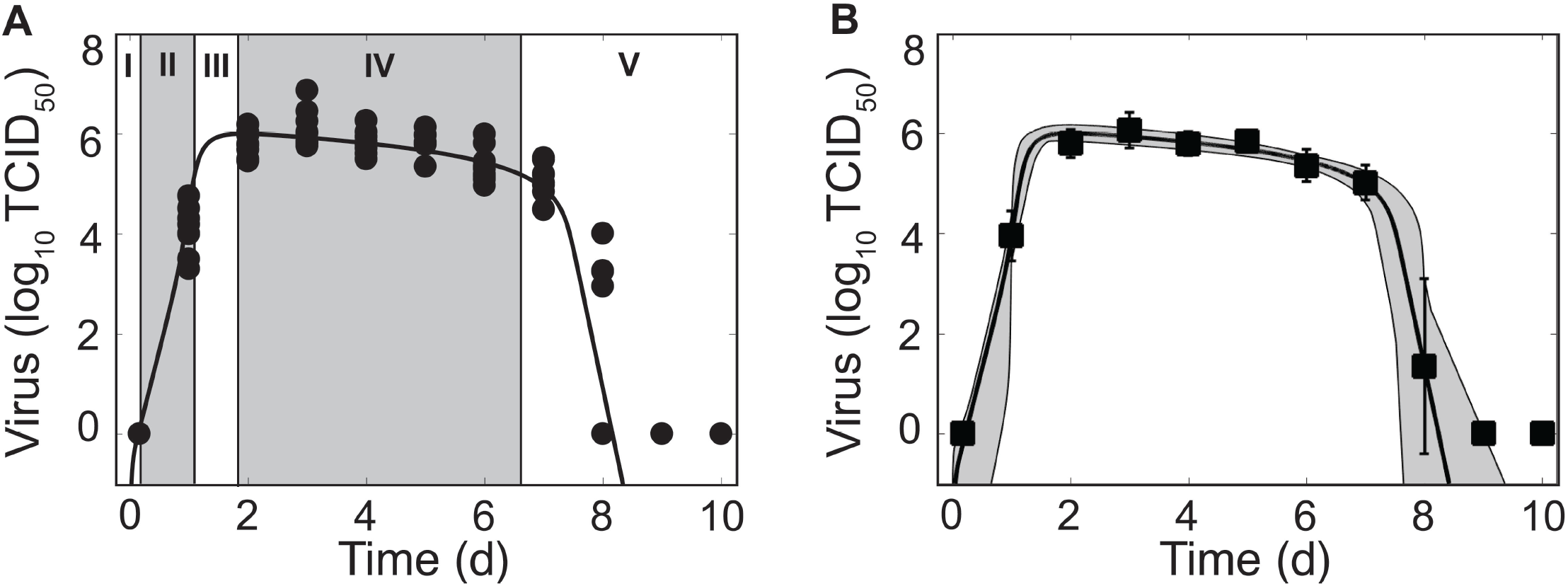
Phases of Virus Kinetics and Fit of the Density-Dependent Viral Kinetic Model. (A) Fit of the density-dependent viral kinetic model (Equations (1)-(5)) to viral lung titers from individual mice (10 mice per time point) infected with 75 TCID_50_ PR8. Each dot is an individual mouse and the solid black line is the optimal solution. Phase I-V of the viral kinetics are shown, where virus (I) quickly infects cells, (II) increases exponentially, (III) peaks, (IV) decays slowly, then (V) decays rapidly and clears. (B) Optimal fit of the model (solid black line) shown with the model solutions using parameter sets within the 95% CIs (gray shading). Data are shown as the mean ± standard deviation. Parameters are given in Table 1.

### Lung Titers

Mice were euthanized by CO_2_ asphyxiation. Lungs were aseptically harvested, washed three times in PBS, and placed in 500 μl sterile PBS. Whole lungs were digested with collagenase (1mg/ml, Sigma C0130), and physically homogenized by syringe plunger against a 40 μm cell strainer. Cell suspensions were centrifuged at 4°C, 500 × g for 7 min. The supernatants were used to determine the viral titers using serial dilutions on MDCK monolayers.

### Mathematical Model

The standard viral kinetic model used to describe IAV infection kinetics tracks 4 populations: susceptible epithelial (“target”) cells (*T*), two classes of infected cells (*I*_1_ and *I*_2_), and virus (*V*) [1]:

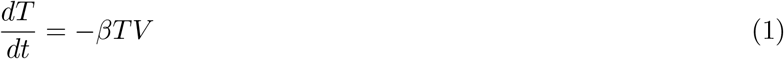

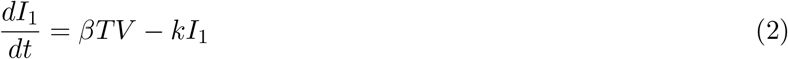

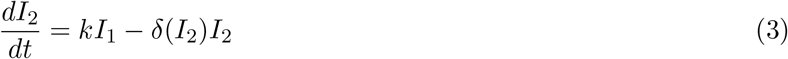

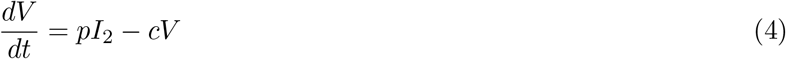

In this model, target cells become infected with virus at rate *βV* per cell. Once infected, these cells enter an eclipse phase (*I*_1_) before transitioning at rate *k* per cell to produce virus (*I*_2_). Virus production occurs at rate *p* per cell. Virus is cleared at rate *c* and virus-producing infected cells (*I*_2_) are cleared according to the function *δ*(*I*_2_). The standard viral kinetic model assumes that infected cells are cleared at a constant rate (*δ*(*I*_2_) = *δ*_*s*_)[1]. The subscript *s* is used to denote “standard”. This model could not recapitulate the data (see Table S1 and Figure S1) and a modification of the model was necessary. Given that the rate of infected cell clearance (*δ*(*I*_2_)) drives the virus decay dynamics [45], we let the clearance rate vary with the number of infected cells such that

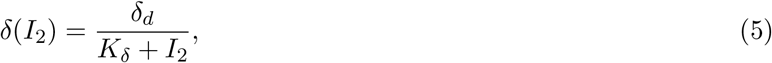

where *δ*_*d*_/*K*_*δ*_ is the maximum rate of infected cell clearance and *K*_*δ*_ is the half-saturation constant. The subscript *d* is used to denote “density-dependent”. Modifications to other terms were examined, but none could replicate the data.

### Parameter Estimation

Given a parameter set *θ*, the cost 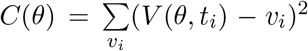 was minimized across parameter ranges using an Adaptive Simulated Annealing (ASA) global optimization algorithm (details in the Supplementary Material) to compare experimental and predicted values of log_10_ TCID_50_/lung. A sample search pattern is shown in Figure S2. Errors of the log_10_ data were assumed to be normally distributed. To explore and visualize the regions of parameters consistent with the models, we fit Equations (1)-(5) to 1000 bootstrap replicates of the data. For each bootstrap data set, the model was fit 10 times beginning from the best-fit parameters estimate *θ*^best^ that was found by fitting the model to the data then perturbing each parameter estimate uniformly within ±50% of its best-fit value. If the three best bootstrap fits were within *χ*^2^ = 0.05 of the best-fit, then the bootstrap was considered successful [46, 48]. For each best fit estimate, we provide 95% confidence interval (CI) computed from the bootstrap replicates. All calculations were performed in MATLAB.

Estimated parameters included the rates of virus infection (*β*), virus production (*p*), virus clearance (*c*), eclipse phase transition (*k*), infected cell clearance (*δ*_*d*_), and the half saturation constant (*K*_*δ*_). Bounds were placed on the parameters to constrain them to physically realistic values. Because biological estimates are not available for all parameters, ranges were set reasonably large based on preliminary results and previous estimates [46]. The rate of infection (*β*) was allowed to vary between 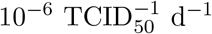 and 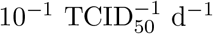, and the rate of viral production (*p*) between 10^−1^ TCID_50_ cell^−1^ d^−1^ and 10^3^ TCID_50_ cell^−1^ d^−1^. Bounds for the viral clearance rate (*c*) were 1 d^−1^ (*t*_1/2_ = 16.7 h) and 10^3^ d^−1^ (*t*_1/2_ = 1 min). Previous estimates of the eclipse phase rate (*k*) for IAV infection in mice resulted in estimates that fell outside the biologically feasible range of 4-6 h [46]. To insure biological feasibility, the lower and upper bounds for the eclipse phase rate (*k*) were 4 d^−1^ and 6 d^−1^. Limits for the half-saturation constant (*K*_*δ*_) were 10^2^ − 10^6^ cells, and limits for the infected cell clearance parameter (*δ*_*d*_) were 1 × 10^6^ − 4 × 10^6^ cells/d.

The initial number of target cells (*T*_0_) was set to 10^7^ cells [46, 48]. Because the initial viral inoculum rapidly infects cells and/or is cleared within 4 h pi, as indicated by the undetectable viral titers at this time point (Figure 1), the initial number of infected cells *I*_1_ (0) was set to 75 cells to reflect an initial dose of 75 TCID_50_. This is an alteration from previous studies, including our own, that either estimate the initial amount of virus (*V*_0_) or set its value to the true viral inoculum. Fixing *V*(0) = 75 TCID_50_ or estimating its value did not improve the fit and could not be statistically justified (see, for example, Table S1). Further, fixing *V*(0) = 75 TCID_50_ yielded an unreasonably high estimate for the rate of virus clearance (*c*) due to the attempt to fit the sharp decay between 0-4 h pi. Estimating *I*_1_(0) could also not be justified and did not improve the model fit (e.g., as in Table S1 and Figure S1). The initial number of productively infected cells (*I*_2_(0)) and the initial free virus (*V*0) were set to 0.

### Linear Regression

We used the function *polyfit* in MATLAB to perform linear regression of the log_10_ values of viral titer during the growth phase (4h, 1 d pi) and the two decay phases (2-6 d pi and 7-8 d pi).

## Results

### Phases of Viral Load Kinetics

Mice infected with 75 TCID_50_ PR8 have viral load kinetics that can be separated into five distinct phases (Figure 1A). This is in contrast to the three phases that we previously defined [45]. In the first phase, virus quickly infects cells and is undetectable within 4 h pi. In the second and third phases, virus increases exponentially and peaks after ~2 d pi. Following the peak, the viral decline can be separated into two phases. In the first decay phase (2-6 d pi), virus decays slowly at a relatively constant rate. In the second decay phase (7-8 d pi), virus declines rapidly (4-5 log_10_ TCID_50_). Sixty percent of mice had no detectable virus by 8 d pi. The remaining mice resolved the infection by 9 d pi.

These data reduced the heterogeneity observed in a previous data set from infection with the same virus [46]. We discovered that the majority of heterogeneity in the previous data set could be attributed to inconsistent infections and, thus, inocula that varied. We further reduced heterogeneity by normalizing the viral titer to the total lung volume, rather than using units of TCID_50_/ml lung homogenate. As expected, some heterogeneity remains at 1 d pi and at 8 d pi. These time points correspond to when virus is rapidly increasing and decreasing, respectively.

### Kinetic Model with Density Dependent Viral Clearance

We first fit the standard viral kinetic model, which is given by Equations (1)-(4) and assumes only one mechanism of constant clearance (*δ*(*I*_2_) = *δ*_*s*_) [1], to the viral load data (see Supplementary Material). This model was unable to capture the entire time course of viral load dynamics, but was able to fit the data from infection initiation to 7 d pi (Figure S1). To more accurately model IAV kinetics and simultaneously recapitulate the two phases of viral decline, we modified the rate of infected cell clearance (*δ*(*I*_2_)) so that the rate changes with respect to the density of the infected cell population. That is, *δ*(*I*_2_) = *δ*_*d*_/(*K*_*δ*_ + *I*_2_) (Equation (5)), where *δ*_*d*_/*K*_*δ*_ is the maximum rate of clearance and *K*_*δ*_ is the number of productively infected cells where the rate is half of its maximum.

Fitting this new model to the viral load data illustrated that the model can accurately reproduce the data and simultaneously capture both phases of viral decline while excluding specific immune responses. The resulting dynamics are shown in Figure 1, the parameter values and 95% confidence intervals (CIs) are given in Table 1, and the parameter ensembles are shown in Figures 2 and S3. For this model, the basic reproduction number (*R*_0_) is given by

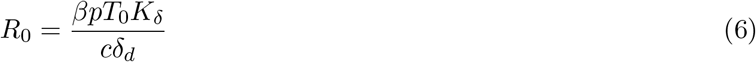

**Figure 2.**
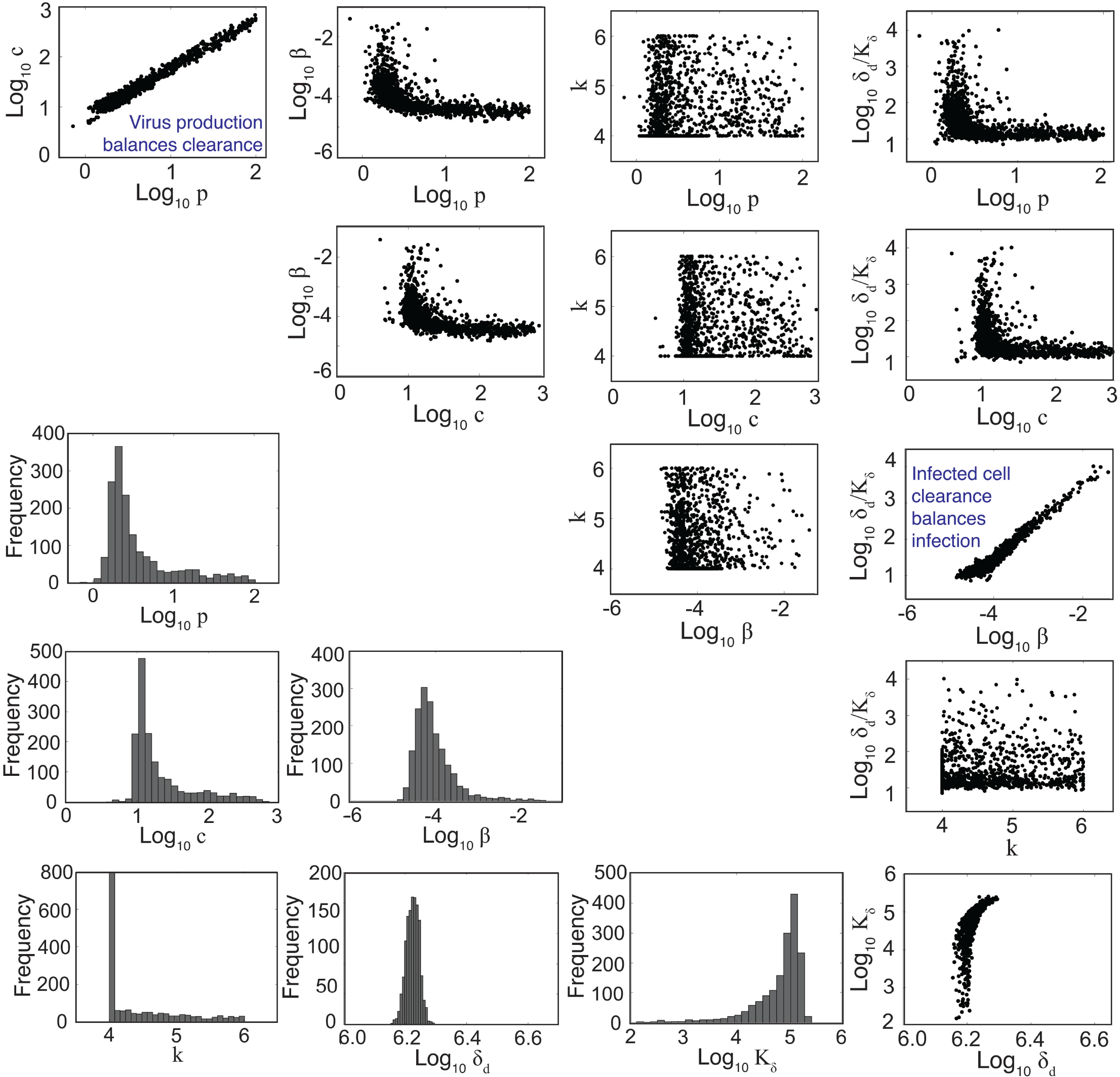
Parameter Ensembles and Histograms. Parameter ensembles and histograms resulting from fitting the density-dependent kinetic model (Equations (1)-(5)) to viral titers from mice infected with 75 TCID_50_ PR8. Two main correlations are evident between the rates of virus production (*p*) and clearance (*c*) and between the rates of infection (*β*) and infected cell clearance (*δ*_*d*_/*K*_*δ*_). The axes limits reflect imposed bounds (except *k* ∈ [4, 6]). All parameters except the eclipse phase rate (*k*) are well bounded (i.e., the 95% CIs do not reach the imposed bounds). Additional ensemble plots (e.g., for *R*_0_) are in Figure S3.

**Table 1.** Parameters and 95% confidence intervals obtained from fitting the density-dependent model (Equations (1)-(5)) to viral titers from mice infected with 75 TCID_50_ PR8.

| Parameter | Description | Units | Value | 95% CI |
| --- | --- | --- | --- | --- |
| $\beta$ | Virus infectivity | TCID<sub>50</sub><sup>-1</sup> d<sup>-1</sup> | $2.4 \times 10^{-4}$ | $[5.0 \times 10^{-5}, 7.8 \times 10^{-2}]$ |
| $p$ | Virus production | TCID<sub>50</sub> cell<sup>-1</sup> d<sup>-1</sup> | 1.6 | [0.82, 125.3] |
| $c$ | Virus clearance | d<sup>-1</sup> | 13.0 | [6.3, 943.1] |
| $k$ | Eclipse phase | d<sup>-1</sup> | 4.0 | [4.0, 6.0] |
| $\delta_d$ | Infected cell clearance | cell<sup>-1</sup> d<sup>-1</sup> | $1.6 \times 10^6$ | $[1.4 \times 10^6, 1.7 \times 10^6]$ |
| $K_\delta$ | Half saturation constant | cells | $4.5 \times 10^5$ | $[1.2 \times 10^2, 1.7 \times 10^5]$ |
| $T(0)$ | Initial uninfected cells | cells | $1 \times 10^7$ | - |
| $I_1(0)$ | Initial infected cells | cells | 75 | - |
| $I_2(0)$ | Initial infected cells | cells | 0 | - |
| $V(0)$ | Initial virus | TCID<sub>50</sub> | 0 | - |

Given the parameters in Table 1, *R*_0_ = 8.8.

To understand how the addition of *δ*(*I*_2_) = *δ*_*d*_/(*K*_*δ*_ + *I*_2_) influences the other parameters during the fitting scheme, we plotted the resulting histograms and 2D parameter projections (Figure 2). As expected, strong correlations exist between the rates of virus production (*p*) and virus clearance (*c*) and between the rate of infection (*β*) and the infected cell death rate (*δ*_*d*_/*K*_*δ*_). The other correlations visible in Figure 2 were a consequence of these two relations. Of note, *δ*_*d*_ was not strongly correlated with any of the other model parameters (Figure S3). In addition, the confidence interval was small, particularly compared to the other parameters. Estimates for the other parameters (*β*, *p*, *c*, and *K*_*δ*_) with the exception of the eclipse phase rate (*k*) were well bounded such that the 95% CIs fell within the upper and lower bounds imposed in the estimation scheme. Similar to previous studies [1, 46], the eclipse phase rate (*k*) restricted to biologically realistic values and was not well defined on the given interval. In support, the ASA algorithm search patterns show a longer search time for *k* compared to the other parameters (Figure S2).

To further determine how the addition of *δ*(*I*_2_) = *δ*_*d*_/(*K*_*δ*_ + *I*_2_) influences the sensitivity of the model solution to changes in parameter values, we performed a one-at-a-time sensitivity analysis (Figure 3). The infected cell clearance parameter (*δ*_*d*_) is the most sensitive parameter and largely dictates the viral decay. Decreasing *δ*_*d*_ significantly delays viral clearance while increasing *δ*_*d*_ leads to rapid viral resolution (Figure 3). In accordance with previous results [45], all other parameters are less sensitive and collectively affect the exponential growth phase and peak.

**Figure 3.**
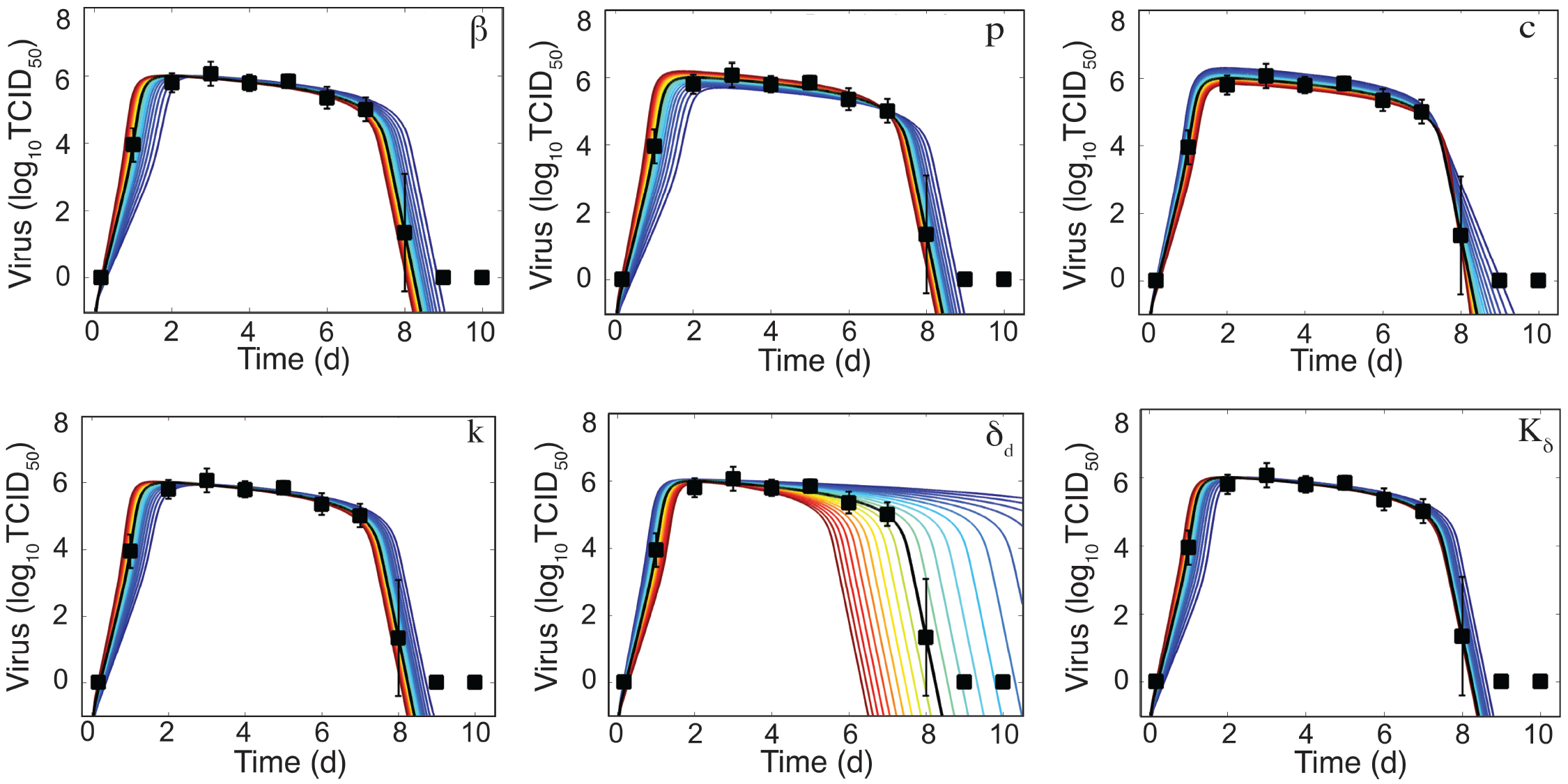
Sensitivity of the Density-Dependent Viral Kinetic Model. Solutions of the density-dependent viral kinetic model (Equations (1)-(5)) for the best-fit parameters (black line, Table 1) and with the indicated parameter increased (red) or decreased (blue) 50% from the best-fit value.

As illustrated in Figure 4A, the rate of infected cell clearance is rapid when these cells are in small numbers. Given the parameters in Table 1, the maximum clearance rate is *δ*(*I*_2_) = 12.7 d^−1^, which corresponds to half-life *t*_1/2_ = 1.3 h. The rate begins to slow when *I*_2_ > 10^4^ cells and is minimal when *I*_2_ is at its maximum (8 × 10^6^ cells). When *I*_2_ is maximal, *δ*(*I*_2_) = 0.21 d^−1^ and *t*_1/2_ = 78 h. In our previous work, we discovered that linear regression analysis could be used to accurately estimate the exponential growth rate, which was a combination of all model parameters, and that the slope of the viral decay could provide an estimate of *δ*(*I*_2_) [45, 46]. To evaluate how these relations correlate to parameters in the model with density dependence, we performed a linear regression on the data during the growth phase (4 h – 1 d pi) and the two decay phases (2–6 d pi and 7–8 d pi) (Figure 4B). The slope of the growth phase is 4.7 log_10_ TCID_50_/d (red line in Figure 4B). In accordance with the previous studies, this slope is a good approximation to the model until shortly before the peak. The model deviates slightly from this estimate and suggests that the virus growth rate briefly increases prior to the peak and that the decay phase begins prior to 2 d pi. This nonlinearity in the growth can be attributed to the decreasing infected cell clearance rate as the number of infected cells increases. These results are in contrast to the standard viral kinetic model, which suggests that the virus growth rate strictly decreases following exponential growth [45, 46]. In the first phase of decay, the slope is −0.2 log_10_ TCID_50_/d (green line in Figure 4B), which corresponds to *δ*(*I*_2_) = 0.4 d^−1^ (green diamond in Figure 4A). In the second phase of decay, the slope is −3.8 log_10_ TCID_50_/d (blue line in Figure 4B), which corresponds to *δ*(*I*_2_) = 8.7 d^−1^ (blue dot in Figure 4A).

**Figure 4.**
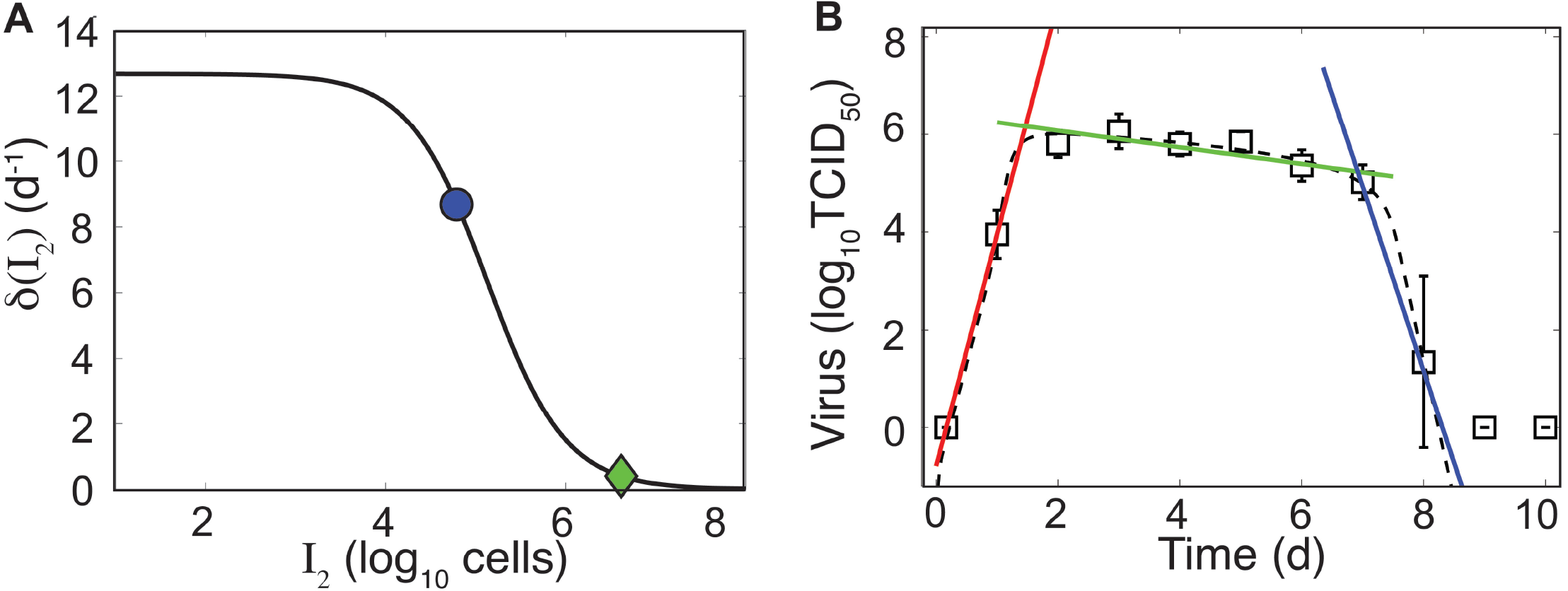
Density-Dependent Infected Cell Clearance Rate and Correlation to Linear Regression. (A) The infected cell clearance rate (*δ*(*I*_2_), Equation (5)) is plotted for different values of infected cells (*I*_2_). The green diamond and the blue dot indicate the corresponding infected cell clearance rates during the slow and fast phases of virus clearance, respectively. These correspond to linear regression estimates in Panel B. (B) Linear regression fits to the viral load data (white squares) during the growth phase (4 h – 1 d pi, red line), the first phase of virus decay (2–6 d pi, green line), or the second phase of virus decay (7–8 d pi, blue line). The dashed black line is the fit of the density-dependent viral kinetic model (Equations (1)–(5)) to the viral load data.

## Discussion

Mathematical models have been widely used to investigate IAV dynamics (reviewed in [3, 7, 41–43]). The viral kinetic model given by Equations (1)-(4) with *δ*(*I*_2_) = *δ*_*s*_ [1] has been the standard in the field for over 10 years. We previously used this model together with data from murine infection to gain insight into IAV virulence factors[46] and into coinfection with bacterial pathogens [40, 44, 48]. Although some predictions made using this model have been experimentally tested and deemed accurate [12, 41, 44, 52], the data here suggested that some dynamical features could not be accounted for and thus a new model was necessary. The model we introduced here includes density-dependent infected cell clearance and better captures the entire course of IAV infection dynamics, including the two-phase viral decay following the peak (Figures 1). Importantly, the model added only a single parameter (the half-saturation constant, *K*_*δ*_) while significantly improving the model fit to viral loads from IAV infection without including additional equations detailing immune responses

By sampling with high frequency and controlling for experimental heterogeneity, we were able to obtain more accurate data (i.e., smaller standard deviations and better reproducibility) that highlighted several important dynamics, some of which were not previously observed. Our data showed that viral loads are maintained at a high level between 2 d and 7 d pi (Figure 1). Sustained viral loads have been observed in several studies [6, 10, 21, 24, 38, 46, 51]. In some data sets, the peak appears more pronounced and is often followed by the plateau phase or a second, lower peak [1, 4, 6, 10, 17, 21, 24]. Our murine data do not indicate a second peak, although there is a subtle increase in viral loads at 5 d pi that may be biologically significant. Previous influenza modeling studies suggest that these dynamics required equations/terms for the innate and adaptive immune responses [1, 9, 33]. However, HIV modeling studies have used similar density-dependent terms to achieve a two-phase viral load decay [8, 20]. Importantly, the model here provides a means for capturing the changes in viral load decay without complicating the model or inferring information about specific immune mechanisms, which are not well understood. However, the change in clearance rate could reflect the change from innate to adaptive immunity. If this is the case, our estimates would suggest that the adaptive response is 25 times more effective than the innate response (−0.2 log_10_ TCID_50_/d between 2–6 d pi versus −3.8 log_10_ TCID_50_/d between 7–8 d pi; Figure 4B).

It is well accepted that the rapid decline in virus during the second decay phase is due to the infiltration of CD8^+^ T cells (reviewed in [13, 23, 27]). These cells typically enter the infection site between 5–6 d pi and peak between 8–9 d pi (e.g., as in [51]). The rapid rate of viral decline between 7–8 d pi suggests that these cells are highly effective. However, the initial infiltration begins at least 1–2 d before a change in the rate of virus decay is visible. Thus, there may be a nonlinearity to this response or it may reflect a simultaneous increase in infections and killing of infected cells or a change in effectiveness proportional to the density of infected cells or to the density of CD8^+^ T cells. A handling-time effect, which can represent the time required for immune cells to kill each infected cell and/or the time for immune cells to become activated (e.g., as in [11, 14, 25, 26, 35, 47]), can slow the per capita rate of clearance. Spatial constraints (e.g., crowding effect), where the number of immune cells within an area is limited, may also play a role. In contrast, numerous clearance mechanisms (e.g., interferons, macrophages, neutrophils, natural killer (NK) cells) are thought to be important during early-and mid-infection, but their contribution to the viral load kinetics is unclear. Using a model to distinguish between these mechanisms is challenging given the close fit of simple kinetic models to viral load data (Figures 1 and S1). Further, neither the data nor the models can discriminate whether the maintenance of high viral loads is due to a lack of clearance of infected cells (i.e., long infected cell lifespan/ineffective clearance) or to the balance of new infections and clearance (i.e., short infected cell lifespan/rapid clearance coupled with rapid virus infection/production). Thus, new experimental designs and more diverse data are necessary.

Viral titers remain the most frequently used data to calibrate models and assess infection dynamics. This is because collecting immunological data is more laborious and expensive. Thus, we seek models that are simple yet accurate and that can be used in the absence of immunological data. The standard viral kinetic model includes the minimal number of parameters and equations needed to recapitulate viral load dynamics. However, viral load data is typically insufficient to uniquely define all 6 parameters [30, 46]. Fortunately, this has not limited our ability to make robust predictions about the underlying biology or to estimate accurate parameter values even when correlations are present [15, 44, 48]. Here, the resulting parameter ensembles were well-bounded (i.e., the 95% CIs did not include the imposed bounds) and correlations were observed in two sets of parameters (Figure 2). The correlation between the rates of virus production (*p*) and virus clearance (*c*) indicates the balance of these processes. This is expected because viral loads measure the amount of virus present and slow virus production/clearance would be indistinguishable from fast production/clearance. Similarly, the rates of infection (*β*) and infected cell clearance (*δ*_*d*_/*K*_*δ*_) were correlated, which indicates a balance of cells becoming infected and being cleared. This is visible in Figure 4B, where the log-linear fit to the data in the growth phase (red line) deviates from the model solution (black dashed line).

Analyzing infection kinetics with mathematical models provides a means to quantify different infection processes. By modeling viral load data, we can make meaningful predictions about the time scales, magnitudes, and rates of different processes even if we cannot directly define specific mechanisms. Further, having a well-characterized model allows us to design new experiments and to perform *in silico* experiments that evaluate situations where data is challenging to obtain. Here, our data, model, and analyses suggest that the clearance rate of infected cells is variable and depends on their density such that clearance slows when infected cells are numerous and is fast when they are in low numbers. Determining what processes give rise to this density dependence remains an open question. Understanding how and why the rate changes should facilitate a deeper understanding of other viral infections and of immunological data, as it becomes available. Further establishing how the virus and host components work together and how they can be manipulated will undoubtedly aid the development of therapies that prevent or treat IAV infections.

## Author Contributions

AMS conceived the idea, performed the experiments, developed the model, ran the simulations, performed the analysis, and wrote the manuscript. APS and DM performed the experiments. VB implemented the code and ran the simulations.

## Acknowledgments

This work was supported by NIH grants AI100946 and AI125324, and by ALSAC. A portion of this work was completed while all authors were at St. Jude Children’s Research Hospital. We thank Alan Perelson and Laura Liao for their helpful comments.

## Supplemental Data

The Supplementary Material includes fits of the standard viral kinetic model, a description of the adaptive simulated annealing (ASA) algorithm, ASA pseudo code, ASA search patterns, and additional parameter ensembles.

